# The phylogenetic placement of an enigmatic moth *Egybolis vaillantina* based on museomics

**DOI:** 10.1101/2022.09.27.509790

**Authors:** Reza Zahiri, Jeremy D. Holloway, Hamid Reza Ghanavi, Leif Aarvik, Niklas Wahlberg

## Abstract

Here, we present multi-locus sequencing results from the enigmatic Afrotropical monotypic genus *Egybolis* Boisduval (occurring in East- and South Africa – previously placed in the subfamily Catocalinae, Noctuidae). Model-based phylogenetic analysis places *Egybolis* within a strongly supported clade comprising four Old World Tropical genera *Cocytia* Boisduval, *Avatha* Walker, *Anereuthina* Hübner and *Serrodes* Guenée from the family Erebidae, subfamily Erebinae. Hence, we propose to formally assign the monotypic genus *Egybolis* to the subfamily Erebinae, and the tribe Cocytiini. Timing of divergence analysis reveals the late Oligocene origin around 25 million years ago (mya) for the tribe Cocytiini, an early Miocene i.e., about 21 mya for the split between *Cocytia* and *Egybolis*.

## Introduction

Megadiverse groups of organisms tend to have taxa with unknown phylogenetic placement, often because they are highly autapomorphic, with a striking appearance, or conversely they lack any shared derived morphological characters with other lineages. Molecular data is finally helping us to sort out the phylogenetic positions of such taxa (e.g. Buckley, Attanayake, and Bradler 2009; Kanda et al. 2016; Sihvonen et al. 2021; Tihelka et al. 2020; Twort et al. 2021), leading to a better understanding of the evolutionary history of these lineages.

Lepidoptera are undoubtedly the largest radiation of phytophagous insects (Scoble, 1992) with over 155,000 described species (Pogue, 2009; van Nieukerken *et al*., 2011). The most diverse lineage of Lepidoptera is undoubtedly the superfamily Noctuoidea with more than 45,890 described species (Pogue, 2009; van Nieukerken *et al*., 2011). Noctuoid species are placed in approximately 5,630 genera (Pogue, 2009) but there are numerous undescribed species, particularly from tropical regions. Moreover, many of these tropical lineages have not been phylogenetically studied thus their evolutionary relationships are unknown. In particular, some of them have been previously assigned to the wrong taxonomic groups (e.g. Sihvonen et al. 2021; Zilli and Grishin 2019).

Not long ago, researchers were using the biological material stored in museum collections only at a morphological study level. Due to the high level of degradation and fragmentation of the genetic material in historical samples, the DNA from these specimens was considered to be too degraded to be utilized in molecular studies (Shapiro and Hofreiter, 2012). Consequently, molecular approaches have been restricted, on one hand, to sequencing technologies (e.g., Sanger sequencing) (Hajibabaei, Singer and Hickey, 2006; Hebert *et al*., 2013), and on the other hand, were limited to quality materials (e.g., freshly collected samples, proper killing agent, etc.). However, recently, next-generation sequencing (NGS) technologies have made the DNA in museum specimens more accessible, either through whole-genome sequencing (WGS) (Sproul and Maddison, 2017; Allio *et al*., 2019; Call *et al*., 2021; Twort *et al*., 2021) or genome reduction methods (Suchan *et al*., 2016; Breinholt *et al*., 2018; Mayer *et al*., 2021). These advanced sequencing approaches have opened up a new field with great potential for studying the evolutionary history of taxa that are difficult to collect: museomics (Call *et al*., 2021).

Here, using a WGS museomics approach, we investigate the phylogenetic position of the enigmatic African genus *Egybolis* Boisduval, 1847 — a monotypic moth genus previously placed in the family Noctuidae. The genus was placed in the subfamily Catocalinae in Noctuidae (e.g. Pinhey, 1975; Vári and Kroon, 1986; Poole, 1989). When the family Erebidae was officially established in 2011 (Zahiri *et al*., 2011) it “automatically” followed the rest of Catocalinae into that family. In the Afromoths website, it is currently listed in Erebidae, Erebinae (De Prins and De Prins, 2011). Its only species, *Egybolis vaillantina* (Stoll, [1790]), the African peach moth, is found in East- and South Africa. Unlike most owlet moths the striking African Peach Moth is diurnal and the larvae (Fig. 1 A, B) feed on peach (Rosaceae) and *Sapindus* (Sapindaceae) species. It has orange antennas and head, a dark blue body, one orange band and two orange spots on each forewing (Fig. 1E, F). Wingspan is 50–60 mm.

**Fig. 1.**
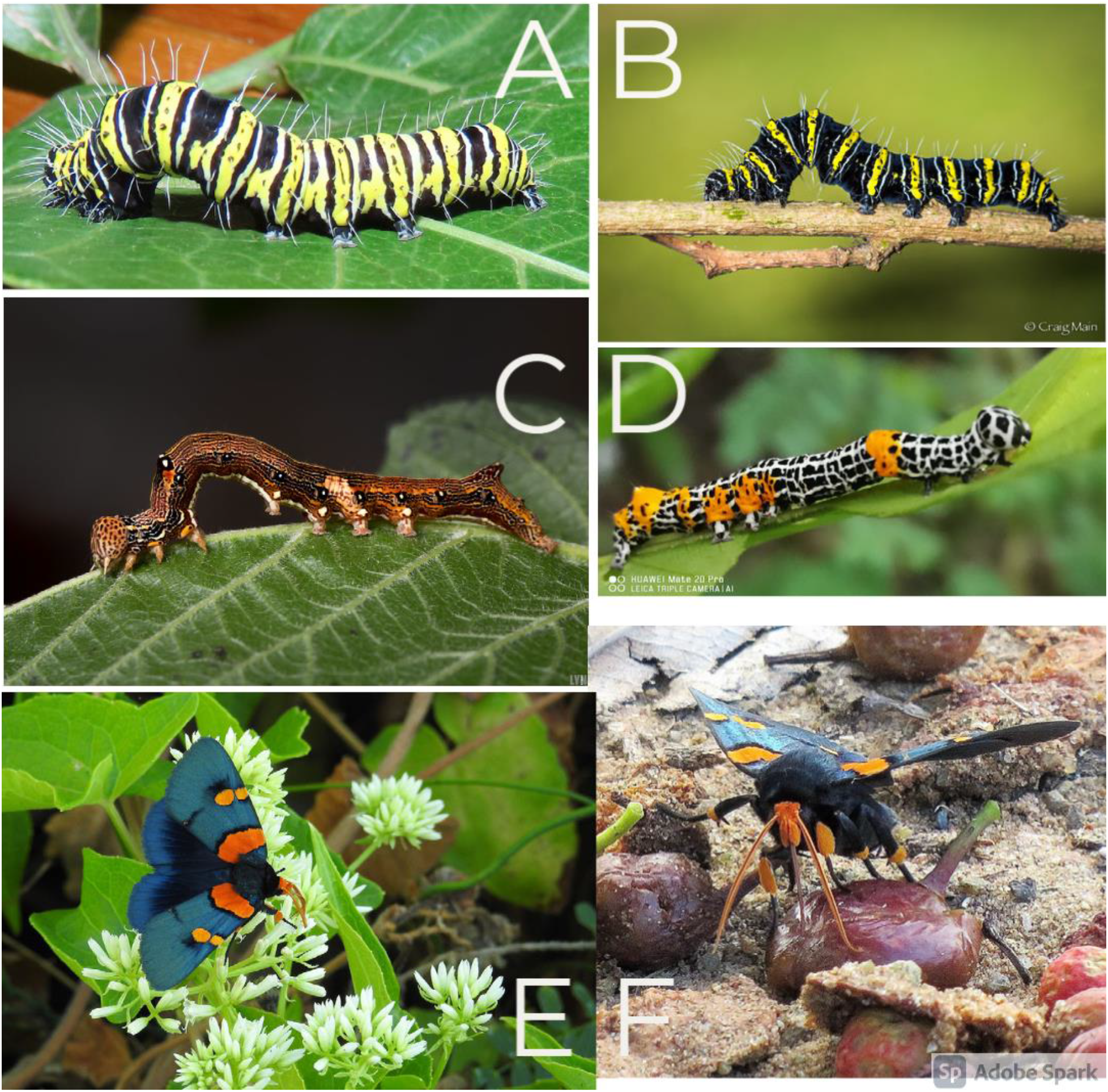
A, B) Caterpillars of the striking African Peach Moth (https://www.inaturalist.org/observations/66703880; https://www.inaturalist.org/observations/48096039); C) Caterpillar of *Avatha* (https://www.inaturalist.org/observations/41976344); D) Caterpillar of *Anereuthina* (https://www.inaturalist.org/observations/95109281); E, F) Adults of the striking African Peach Moth (https://www.inaturalist.org/observations/84578502; https://www.inaturalist.org/observations/67839106).

## Material and methods

Genomic DNA was extracted from a leg of an individual collected in 1993 in Tanzania, currently housed at the Natural History Museum, University of Oslo. The extraction was done in 2011 with the original intention to sequence PCR products from the specimen (according to protocols described in Zahiri et al. (2012). However, all PCRs failed and the extract has been stored at +2°C since then. Here we use the same extract for whole genome shotgun sequencing following protocols published in Twort et al. (2021). Briefly, DNA was blunt-end repaired with T4 Polynucleotide Kinase (New England Biolabs), followed by a reaction clean-up with the MinElute purification kit (Qiagen). This was followed by adapter ligation, reaction purification, and adapter fill-in. The resulting reactions were then indexed using unique dual indexes. Indexing PCR was carried out in six independent reactions to avoid amplification bias, with 15 cycles for each reaction. Indexing PCR reactions were pooled prior to magnetic bead clean-up with Sera-Mag SpeedBeads ™ carboxylate-modified hydrophilic (Sigma-Aldrich). An initial bead concentration of 0.5X was used to remove long fragments that are likely to represent contamination from fresh DNA and libraries were selected with a bead concentration of 1.8X to size select the expected library range of ~300 bp. The resulting library was quantified and quality checked with Quanti-iTTM PicoGreenTM dsDNA assay and with Bioanalyzer 2100, respectively. The final indexed library was pooled with 53 other samples prior to sequencing on an Illumina Novaseq platform (one lane, 2×150 bp, S4 flow cell) at Swedish National Genomics Infrastructure (NGI) in Stockholm.

Following Twort et al. (2021), we checked the quality of the raw reads with FASTQC v0.11.5 (Andrews, 2010). We removed reads with ambiguous bases (N’s) from the dataset using Prinseq 0.20.4 (Schmieder and Edwards, 2011). We then used Trimmomatic 0.38 (Bolger, Lohse, and Usadel, 2014) to remove low-quality bases from their beginning (LEADING: 3) and end (TRAILING: 3), using a sliding window approach. Quality was measured for sliding windows of four base pairs and had to be higher than 25 on average. Reads below 30 bp were also removed. We then de novo assembled the genome of *Egybolis* with SPAdes v3.13.1 (Bankevich *et al*., 2012) with k-mer values of 21, 33, and 55.

We extracted the standard eight gene regions used by Zahiri et al. (2012) using TBLASTN, with *Bombyx mori* amino acid sequences as the reference, as all of these sequences represent one exon within the gene of interest. A preliminary analysis using the dataset published by Zahiri et al. (2011) clearly placed *Egybolis* within Erebidae, thus we used the dataset of Zahiri et al. (2012) for further analyses.

Model-based phylogenetic analyses were based on DNA sequences of eight protein-coding genes — cytochrome *c* oxidase subunit I (COI) from the mitochondrial genome, and elongation factor-1α (EF-1α), ribosomal protein S5 (RpS5), carbamoylphosphate synthase domain protein (CAD), cytosolic malate dehydrogenase (MDH), glyceraldehyde-3-phosphate dehydrogenase (GAPDH), isocitrate dehydrogenase (IDH) and wingless genes from the nuclear genome — from 201 representatives of the major lineages of Erebidae and 36 representatives from the five other families of Noctuoidea: Oenosandridae (one species), Notodontidae (three species), Nolidae (five species), Euteliidae (four species) and Noctuidae (23 species), giving a total of 237+1 (= *Egybolis vaillantina*) terminal taxa (dataset A, Table S1). We included an additional species from BOLD (Ratnasingham and Hebert, 2007) (with only barcode region data, 658 bp) *Anereuthina renosa* (representing type species of the genus) — a genus that appears to be closely related to the *Serrodes* Guenée group (Holloway, 2005).

We ran maximum likelihood (ML) analyses with the dataset partitioned by gene using IQ-TREE 2.0.6 (Trifinopoulos et al. 2016). The best-fitting substitution models and optimal number of partitions were selected by ModelFinder (-m MFP+MERGE) (Kalyaanamoorthy *et al*., 2017). Support for nodes was evaluated with 1000 ultrafast bootstrap (UFBoot2) approximations (Hoang *et al*., 2018), and SH-like approximate likelihood ratio test (Guindon *et al*., 2010) using “-B 1000 -alrt 1000” options. SH-Like ≥80 and UFBoot2 ≥ 95 values indicate well-supported clades. To reduce the risk of overestimating branch supports in UFBoot2 test, we used the -bnni option, which optimizes each bootstrap tree using a hill-climbing nearest neighbour interchange (NNI) search. We rooted the cladograms on *Oenosandra boisduvalii* Newman, which is the putative sister family to the remainder of the Noctuoidea (Regier *et al*., 2009; Mutanen, Wahlberg and Kaila, 2010; R. Zahiri *et al*., 2011; Zahiri *et al*., 2012).

Finally, divergence times were estimated using a relaxed lognormal molecular clock model as implemented in BEAST 2.6.3 (Bouckaert *et al*., 2019). We applied a reduced dataset with 209 terminals (dataset B, Table S2) to estimate the dates of cladogenetic events. We assume that different positions in the alignment could potentially accumulate substitutions differently. As a result, we unlinked the partitions for the substitution models, allowing each partition to evolve under a different substitution model, but we linked the partitions for the clock and tree models. The best partioning scheme and their substitution models were selected using ModelFinder implemented in IQ-TREE (-m TESTMERGEONLY -mset mrbayes), to choose the best set of substitution models in BEAST. The tree prior was set to a Birth Death model. Lepidoptera are characterized by a lack of fossils that can be confidently assigned to extant clades (Grimaldi and Engel, 2005; Wahlberg *et al*., 2013), and noctuoid moths are no exception. As a result, we relied on calibrations derived from phylogenomic study of Kawahara et al. (2019) using 2000 orthologous protein-coding genes of representatives of nearly all lepidopteran superfamilies, and multiple fossil calibrations. We constrained the root of the superfamily Noctuoidea (69 my) with normal distributions encompassing the 95% credibility intervals estimated in Kawahara et al. (2019). Then we constrained the roots of the family Erebidae and the subfamily Arctiinae to 47.8 my and 30 my, respectively, with normal distributions encompassing the 95% credibility intervals (see Dataset S11 Kawahara et al. 2019). To make the run computationally easier, we used phylogenetic constraints, forcing the monophyly of the well supported clades in our phylogenetic results. For a list of the constraints check the Dataset S1.XML file in the supplementary material. The MCMC parameters were fixed to 100 million generations with sampling every 5,000 generations and the first 25% discarded as burn-in. We ran the analysis four independent times (different seed values) and confirmed the convergence of the runs before combining the results in LogCombiner (part of BEAST2 package). We used TreeAnnotator (part of BEAST2 package) to generate the maximum clade credibility (MCC) tree, representing the mean and 95% HPD interval for all nodal ages.

Distribution data for the target species and its closely related species (tribe Cocytiini) was acquired from the iNaturalist API using their tools and georeferenced in QGIS v.3.20.3 (QGIS Development Team, 2022) — a free and open source Geographic Information System — to visualize species distributions.

## Results

We were able to recover sequences of seven of the eight gene regions, only wingless was not recovered from the genome assembly of *Egybolis*. The optimal topology found by dataset A in our analysis and those of the Erebidae dataset of Zahiri et al. (2012) are very similar with slight differences in a few terminal taxa placements. Our results fully resolve the phylogenetic position of the enigmatic *Egybolis* moth with confidence and suggested a close relationship (SH-Like = 100 / aBayes = 1 / UFBoot2 = 100) within the tribe Cocytiini Boisduval, 1829 (Fig. 2). *Egybolis* is placed within the Erebinae in a strongly-supported sister relationship with the genera of the *Serrodes* group of Holloway (2005) and the striking New Guinea and Moluccan genus *Cocytia* in Cocytiini (Zahiri et al. 2012) (Fig. 2). The three genera (i.e., *Serrodes, Avatha*, and *Anereuthina*) included by Holloway (2005) shared general similarity of facies, particularly the presence and disposition of blocks of black on the forewing.

**Fig. 2.**
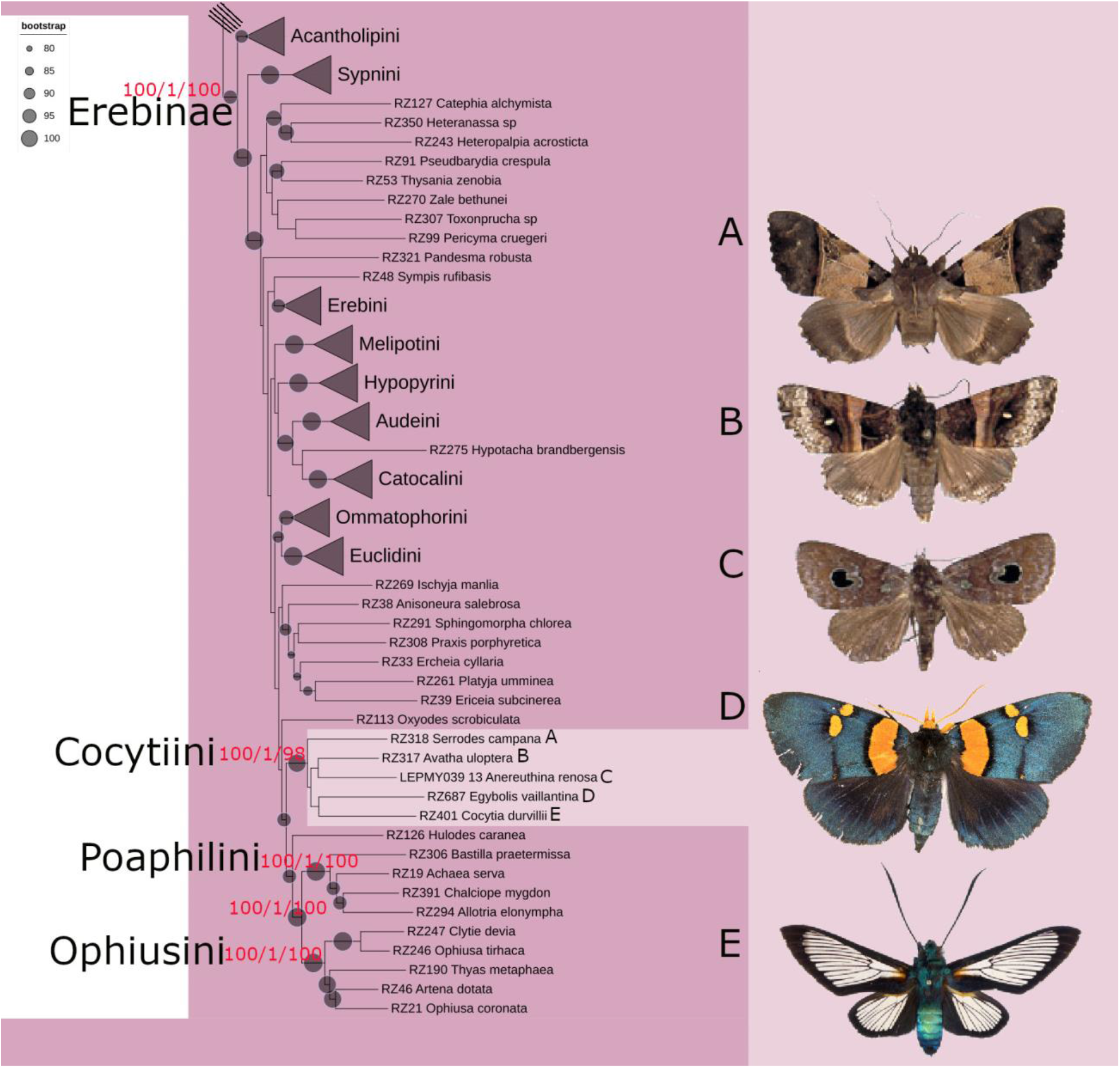
The Maximum likelihood tree from analysis of the eight-gene regions of Erebidae dataset of Zahiri et al. (2012) and additional terminals. The subfamily Erebinae is only shown here (see suppl. Fig 1 for a full tree). Ultrafast bootstrap values (please see legend) are displayed as grey circles on the tree branches. Red numbers above the selected nodes (Erebinae, Cocytiini, Poaphilini, Ophiusini, Poaphilini+Ophiusini) are SH-aLRT support (%)/aBayes support/ultrafast bootstrap support (%). Names of moths shown in the figure from top to bottom: A) *Serrodes campana*, B) *Avatha uloptera*, C) *Anereuthina renosa*, D) *Egybolis vaillantina* and E) *Cocytia durvillii*.

The results of this study corroborate those of Zahiri et al. (2012) by confirming a strongly supported sister relationship between Cocytiini and a clade containing the tribes Poaphilini + Ophiusini (Fig. 2). This clade is well supported (SH-Like = 99.9/aBayes = 1/UFBoot2 = 100) and is divided into two major assemblages, tribes Ophiusini and Poaphilini, each with strong support for reciprocal monophyly (SH-Like = 100/aBayes = 1/UFBoot2 = 100 (Fig. 2).

The chronogram (Fig. 3) obtained from the Bayesian dating analyses allows us to infer a median age of 24.96 myr (late Oligocene) for the tribe Cocytiini, with 95% highest posterior densities (HPD) of 20.49–29.34 myr. Within Cocytiini, *Serrodes* is the first lineage to branch off followed by *Avatha* (about 23.39 mya) and finally with the bifurcation of *Egybolis* from *Cocytia* happening in the late Miocene some 21.12 mya (Fig. 3). Our analysis of divergence time suggests that the ancestors of Erebinae began diversifying in the mid-Eocene some 46.33 mya (Fig. 3).

**Fig. 3.**
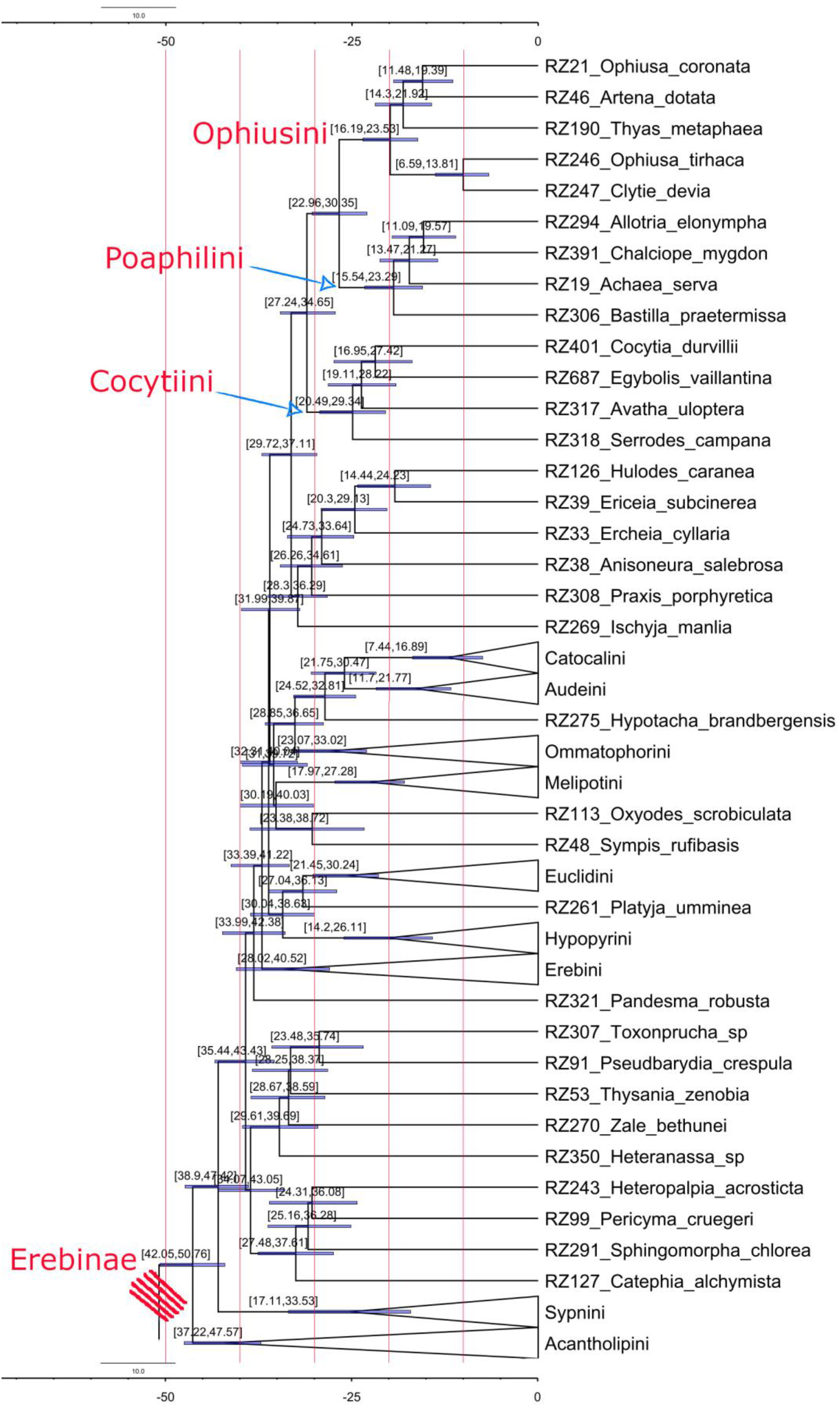
Maximum credibility tree with median ages (myr) from the Bayesian uncorrelated uniform analysis under BEAST for the tribe Cocytiini. A ten Myr-timescale is placed at the top of the chronogram. Horizontal blue bands represent the 95% HPD heights (in myr) for the major nodes of the chronogram. Node numbers in brackets above the nodes are the 95% HPD heights (in myr).

## Discussion

Our study offered an eminent example of sharing strong phylogenetic signals and connections among the geographically separated organisms in the Austral Region. We find that closely related members of Cocytiini moths are widely separated on the various austral landmasses (Fig. 4). The enigmatic Afrotropical *Egybolis* moth has a close evolutionary relationship with other members of the tribe Cocytiini (Erebidae, Erebinae) that are principally distributed in humid climates in tropical regions of the Old World. *Avatha* is Indo-Malayan/Australasian genus, *Serrodes* is an Old-Tropic genus (Indo-Malayan/Australasian/African), *Anereuthina* is Indo-Malayan, and *Cocytia* is New Guinea and Moluccan genus (Fig. 4).

**Fig. 4.**
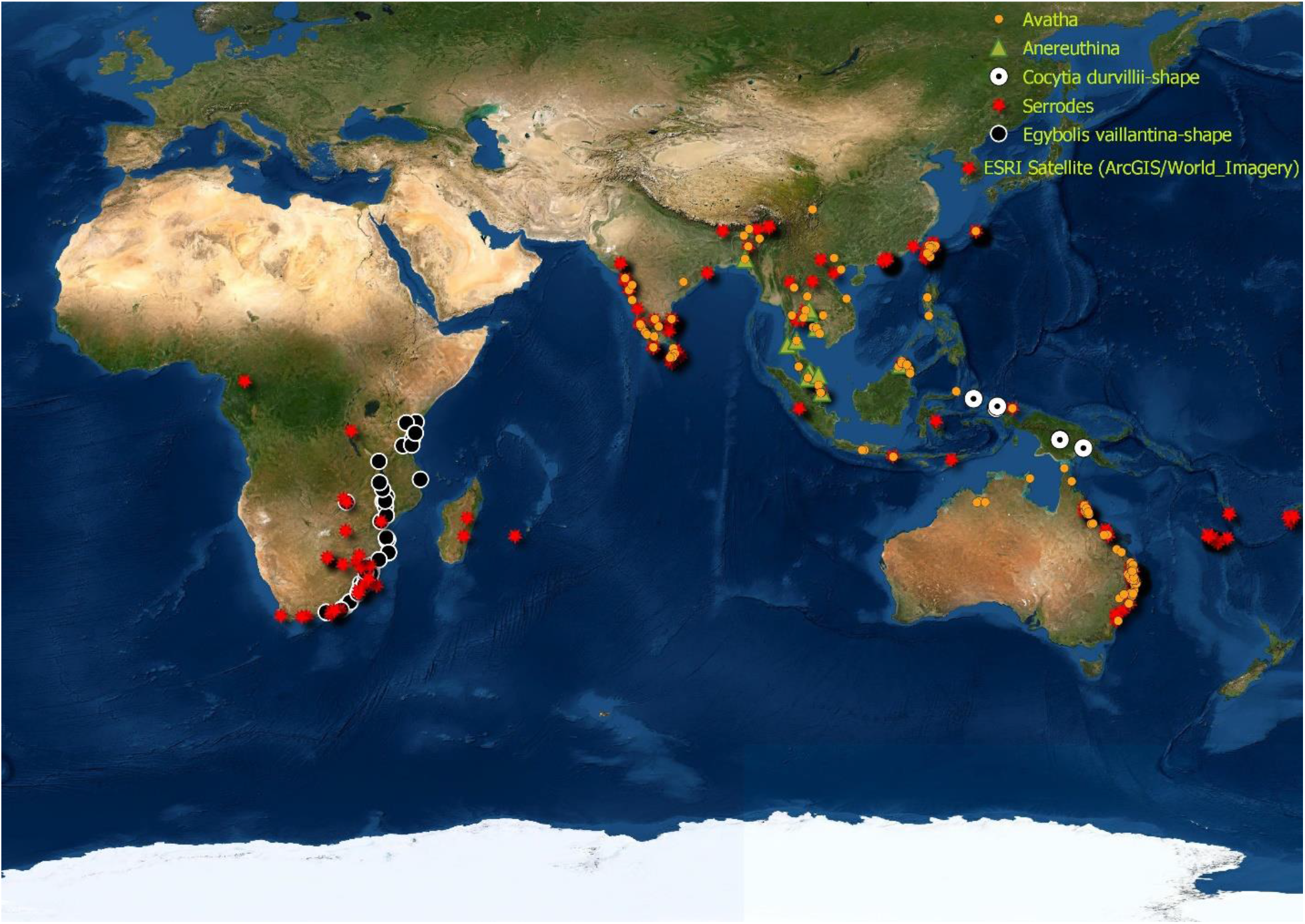
Distribution map of members of the tribe Cocytiini (Erebidae, Erebinae).

*Cocytia* typifies the tribe Cocytiini and was illustrated in Holloway et al. (2001). The status of *Cocytia* was also studied by Speidel et al. (1997), with illustrations, and further details of its morphology presented by Heppner (2010). Speidel et al. (1997) noted that the proboscis has a smooth, rather than nodulose, apex, and all the styloconic sensilla are dorsal. This condition is shared with many genera that they examined in Erebinae as constituted in the study of Zahiri et al. (2012). The structures of the male genitalia could also be interpreted as being a modification of the condition in *Avatha* (see Holloway 2005). The larvae of the *Serrodes* group of Holloway are of the ophiusine type, and a pupal bloom has been noted in a species of *Avatha* Walker, but not in *Serrodes* or *Anereuthina*. The prolegs all appear well developed except those on A3 which are reduced; the plantae are expanded to give each a T-shape. The condition of reduced prolegs on A3 is exactly the same condition that occurs in the larvae of *Egybolis* (Fig. 1). Host plants in *Serrodes* and *Avatha* are predominantly from the Sapindaceae, but *Anereuthina* larvae are palm-feeders. The larvae of *Egybolis* feed on peach and *Sapindus* species. The strong association from DNA sequence evidence and morphology of the members of Cocytiini led Zahiri et al. (2012) to suggest that the larva of *Cocytia* may also feed on Sapindaceae, like those of *Serrodes* and *Avatha* as well as *Egybolis* in this study. *Avatha* includes Indo-Australian species (Fig. 3) of the old but inappropriate concept of *Athyrma* Hübner (type species *adjutrix* Cramer, Surinam), a genus essentially restricted to South America and not related to the species discussed here. *Serrodes* species are Afro-Indo-Australian, and *Anereuthina* includes Indo-Malayan species (Fig. 3). *Egybolis* and *Cocytia* are both monotypic genera with restricted geographical distributions —found in southeastern Africa and New Guinea, respectively—compared to their sister taxa (Fig. 3).

The diversification of the tribe Cocytiini in the Miocene is similar to other groups of Lepidoptera that have connections between Africa and Southeast Asia (e.g. Kodandaramaiah and Wahlberg 2007; Aduse-Poku et al. 2009; Kawahara et al. 2019; Toussaint et al. 2019; Chazot et al. 2021). The sister relationship of *Egybolis* and *Cocytia*, and their disjunct distributions in Africa and New Guinea, respectively, suggest that their common ancestor may have been widespread from Africa to New Guinea. Indeed, there may be extant species that are found currently in Southeast Asia that are more related to one or the other, that we simply have not yet investigated. Given the surprising phylogenetic position of *Egybolis*, this is quite a likely scenario.

## Conclusion

In conclusion, using a museomics approach, we have been able to ascertain the phylogenetic position of an enigmatic moth species, and thus transfer *Egybolis vaillantina* from Noctuidae to Erebidae: Erebinae: Cocytiini. Museomics approaches show great potential for resolving such mysteries, which megadiverse groups of organisms tend to be laden with. Given that the whole genome of *Egybolis* was sequenced in this study, the data can be reused in the future in genome-level studies, including phylogenomic analyses of Erebidae (e.g. Homziak et al. 2019; Ghanavi et al. 2022).

## Acknowledgments

This work was financially supported by the Swedish Research Council (Grant No. 2015-04441) awarded to N. Wahlberg; CIMO + Finnish Cultural Foundation + Alfred Kordelin Foundation awarded to R. Zahiri.

## Supporting Information

**Table S1.** List of taxa with voucher codes (specimen ID = specimen identity) and GenBank accession numbers used in the multi-locus analysis. - = Gene was not amplified for specimens. X = to be submitted to GenBank to acquire accession numbers.

**Table S2.** List of taxa with voucher codes (specimen ID = specimen identity) and GenBank accession numbers used in time of divergences analysis. - = Gene was not amplified for specimens. X = to be submitted to GenBank to acquire accession numbers.

**Supplementary Fig. 1.** The Maximum likelihood tree from analysis of the eight-gene regions of Erebidae dataset of Zahiri et al. (2012) and additional terminals. Ultrafast bootstrap values (please see legend) are displayed as grey circles on the tree branches.

**Dataset S1.** XML input file for BEAST generated in BEAUti (part of BEAST2 package).

## References

Aduse-Poku, K., Vingerhoedt, E. and Wahlberg, N. (2009) ‘Out-of-Africa again: a phylogenetic hypothesis of the genus Charaxes (Lepidoptera: Nymphalidae) based on five gene regions.’, Molecular phylogenetics and evolution, 53(2), pp. 463–478. Available at: https://doi.org/10.1016/j.ympev.2009.06.021.

Allio, R. et al. (2019) ‘Whole genome shotgun phylogenomics resolves the pattern and timing of swallowtail butterfly evolution’, Systematic Biology [Preprint]. Available at: https://doi.org/10.1093/sysbio/syz030.

Andrews, S. (2010) Babraham Bioinformatics - FastQC A Quality Control tool for High Throughput Sequence Data. Available at: https://www.bioinformatics.babraham.ac.uk/projects/fastqc/ (Accessed: 24 August 2022).

Bankevich, A. et al. (2012) ‘SPAdes: a new genome assembly algorithm and its applications to single-cell sequencing.’, Journal of computational biology : a journal of computational molecular cell biology, 19(5), pp. 455–477. Available at: https://doi.org/10.1089/cmb.2012.0021.

Bolger, A.M., Lohse, M. and Usadel, B. (2014) ‘Trimmomatic: a flexible trimmer for Illumina sequence data’, Bioinformatics, 30(15), pp. 2114–2120. Available at: https://doi.org/10.1093/bioinformatics/btu170.

Bouckaert, R. et al. (2019) ‘BEAST 2.5: An advanced software platform for Bayesian evolutionary analysis’, PLOS Computational Biology, 15(4), p. e1006650. Available at: https://doi.org/10.1371/journal.pcbi.1006650.

Breinholt, J.W. et al. (2018) ‘Resolving Relationships among the Megadiverse Butterflies and Moths with a Novel Pipeline for Anchored Phylogenomics’, Systematic Biology, 67(1), pp. 78–93. Available at: https://doi.org/10.1093/sysbio/syx048.

Buckley, T.R., Attanayake, D. and Bradler, S. (2009) ‘Extreme convergence in stick insect evolution: phylogenetic placement of the Lord Howe Island tree lobster’, Proceedings of the Royal Society B: Biological Sciences, 276(1659), pp. 1055–1062. Available at: https://doi.org/10.1098/rspb.2008.1552.

Call, E. et al. (2021) ‘Museomics: Phylogenomics of the Moth Family Epicopeiidae (Lepidoptera) Using Target Enrichment’, Insect Systematics and Diversity, 5(2), p. 6. Available at: https://doi.org/10.1093/isd/ixaa021.

Chazot, N. et al. (2021) ‘Conserved ancestral tropical niche but different continental histories explain the latitudinal diversity gradient in brush-footed butterflies’, Nature Communications, 12(1), p. 5717. Available at: https://doi.org/10.1038/s41467-021-25906-8.

De Prins, J. and De Prins, W. (2011) Afromoths, online database of Afrotropical moth species (Lepidoptera). World Wide Web electronic publication (http://www.afromoths.net) [2022-08-04].Available at: http://www.afromoths.net/species/show/34593 (Accessed: 4 August 2022).

Ghanavi, H.R. et al. (2022) ‘The (non) accuracy of mitochondrial genomes for family-level phylogenetics in Erebidae (Lepidoptera)’, Zoologica Scripta, n/a(n/a). Available at: https://doi.org/10.1111/zsc.12559.

Grimaldi, D. and Engel, M.S. (2005) Evolution of the insects. Cambridge, New York etc.: Cambridge University Press.

Guindon, S. et al. (2010) ‘New algorithms and methods to estimate maximum-likelihood phylogenies: Assessing the performance of PhyML 3.0’, Systematic Biology, 59(3), pp. 307–321. Available at: https://doi.org/10.1093/sysbio/syq010.

Hajibabaei, M., Singer, G.A. and Hickey, D.A. (2006) ‘Benchmarking DNA barcodes: an assessment using available primate sequences’, Genome, 49(7), pp. 851–854. Available at: https://doi.org/10.1139/g06-025.

Hebert, P.D.N. et al. (2013) ‘A DNA “Barcode Blitz”: Rapid Digitization and Sequencing of a Natural History Collection’, PloS one, 8(7), p. e68535. Available at: https://doi.org/10.1371/journal.pone.0068535.

Heppner, J.B. (2010) ‘Notes on the Moluccan and Papuan genus Cocytia (Lepidoptera: Noctuidae: Cocytiinae)’, Lepidoptera Novae, 3(4), pp. 217–221.

Hoang, D.T. et al. (2018) ‘UFBoot2: Improving the Ultrafast Bootstrap Approximation’, Molecular Biology and Evolution, 35(2), pp. 518–522. Available at: https://doi.org/10.1093/molbev/msx281.

Holloway, J.D. (2001) ‘The Moths of Borneo (part 7): Family Arctiidae, subfamily Lithosiinae’, Malayan Nature Journal, 55, pp. 279–458.

Holloway, J.D. (2005) ‘The Moths of Borneo (part 15 & 16): Family Noctuidae, subfamily Catocalinae’, Malayan Nature Journal, 58, pp. 1–529.

Homziak, N.T. et al. (2019) ‘Anchored hybrid enrichment phylogenomics resolves the backbone of erebine moths’, Molecular Phylogenetics and Evolution, 131, pp. 99–105. Available at: https://doi.org/10.1016/j.ympev.2018.10.038.

Kalyaanamoorthy, S. et al. (2017) ‘ModelFinder: Fast model selection for accurate phylogenetic estimates’, Nature Methods [Preprint]. Available at: https://doi.org/10.1038/nmeth.4285.

Kanda, K. et al. (2016) ‘Phylogenetic placement of the Pacific Northwest subterranean endemic diving beetle Stygoporus oregonensis Larson & LaBonte (Dytiscidae, Hydroporinae)’, ZooKeys, 632, pp. 75–91. Available at: https://doi.org/10.3897/zookeys.632.9866.

Kawahara, A.Y. et al. (2019) ‘Phylogenomics reveals the evolutionary timing and pattern of butterflies and moths’, Proceedings of the National Academy of Sciences, 116(45), p. 22657. Available at: https://doi.org/10.1073/pnas.1907847116.

Kodandaramaiah, U. and Wahlberg, N. (2007) ‘Out-of-Africa origin and dispersal-mediated diversification of the butterfly genus Junonia (Nymphalidae: Nymphalinae)’, Journal of Evolutionary Biology, 20(6), pp. 2181–2191. Available at: https://doi.org/10.1111/j.1420-9101.2007.01425.x.

Mayer, C. et al. (2021) ‘Adding leaves to the Lepidoptera phylogeny: Capturing hundreds of nuclear genes from old museums specimens’, Systematic Entomology [Preprint]. Available at: https://doi.org/DOI:10.1111/syen.12481.

Mutanen, M., Wahlberg, N. and Kaila, L. (2010) ‘Comprehensive gene and taxon coverage elucidates radiation patterns in moths and butterflies’, Proceedings of the Royal Society B-Biological Sciences, 277(1695), pp. 2839–2848.

Nguyen, L.-T. et al. (2015) ‘IQ-TREE: A Fast and Effective Stochastic Algorithm for Estimating Maximum-Likelihood Phylogenies’, Molecular Biology and Evolution, 32(1), pp. 268–274. Available at: https://doi.org/10.1093/molbev/msu300.

van Nieukerken, E.J. et al. (2011) ‘Order Lepidoptera Linnaeus, 1758.’, Zootaxa, 3148, pp. 212–221.

Pinhey, E.C.G. (1975) Moths of Southern Africa. Cape Town, South Africa: Tafelberg Publishers.

Pogue, M.G. (2009) ‘Lepidoptera Biodiversity’, in R.G. Foottit and P.H. Adler (eds) Insect Biodiversity: Science and Society. Wiley-Blackwell.

Poole, R.W. (1989) Lepidopterorum Catalogus (New Series). Fascicle 118. Noctuidae. New York: E.J. Brill/Flora and Fauna Publications.

QGIS Development Team (2022) ‘QGIS Geographic Information System. Open Source Geospatial Foundation Project.’ Available at: http://qgis.osgeo.org.

Ratnasingham, S. and Hebert, P.D.N. (2007) ‘BARCODING: bold: The Barcode of Life Data System (http://www.barcodinglife.org): BARCODING’, Molecular Ecology Notes, 7(3), pp. 355–364. Available at: https://doi.org/10.1111/j.1471-8286.2007.01678.x.

Regier, J.C. et al. (2009) ‘Toward reconstructing the evolution of advanced moths and butterflies (Lepidoptera: Ditrysia): an initial molecular study’, Bmc Evolutionary Biology, 9.

Schmieder, R. and Edwards, R. (2011) ‘Quality control and preprocessing of metagenomic datasets.’, Bioinformatics (Oxford, England), 27(6), pp. 863–864. Available at: https://doi.org/10.1093/bioinformatics/btr026.

Scoble, M.J. (1992) The lepidoptera: form, function, and diversity. Oxford;New York; Oxford University Press (Book, Whole).

Shapiro, B. and Hofreiter, M. (2012) Ancient DNA. Methods and Protocols. Springer New York Dordrecht Heidelberg London: Humana Press.

Sihvonen, P. et al. (2021) ‘Insect taxonomy can be difficult: a noctuid moth (Agaristinae: Aletopus imperialis) and a geometrid moth (Sterrhinae: Cartaletis dargei) combined into a cryptic species complex in eastern Africa (Lepidoptera).’, PeerJ, 9, p. e11613. Available at: https://doi.org/10.7717/peerj.11613.

Speidel, W., Fanger, H. and Naumann, C.M. (1997) ‘On the systematic position of Cocytia Boisduval, 1828 (Lepidoptera:Noctuidae)’, Deutsche Entomologische Zeitschrift, 44(1), pp. 27–31.

Sproul, J.S. and Maddison, D.R. (2017) ‘Sequencing historical specimens: successful preparation of small specimens with low amounts of degraded DNA’, Molecular Ecology Resources, 17(6), pp. 1183–1201. Available at: https://doi.org/10.1111/1755-0998.12660.

Suchan, T. et al. (2016) ‘Hybridization Capture Using RAD Probes (hyRAD), a New Tool for Performing Genomic Analyses on Collection Specimens’, PLOS ONE, 11(3), p. e0151651. Available at: https://doi.org/10.1371/journal.pone.0151651.

Tihelka, E. et al. (2020) ‘Fleas are parasitic scorpionflies’, Palaeoentomology, 3(6), pp. 641–653. Available at: https://doi.org/10.11646/palaeoentomology.3.6.16.

Toussaint, E.F.A. et al. (2019) ‘Out of the Orient: Post-Tethyan transoceanic and trans-Arabian routes fostered the spread of Baorini skippers in the Afrotropics’, Systematic Entomology, 44(4), pp. 926–938. Available at: https://doi.org/10.1111/syen.12365.

Trifinopoulos, J. et al. (2016) ‘W-IQ-TREE: a fast online phylogenetic tool for maximum likelihood analysis’, Nucleic acids research, 44(W1), pp. W232–W235. Available at: https://doi.org/10.1093/nar/gkw256.

Twort, V.G. et al. (2021) ‘Museomics of a rare taxon: placing Whalleyanidae in the Lepidoptera Tree of Life’, Systematic Entomology, 46(4), pp. 926–937. Available at: https://doi.org/10.1111/syen.12503.

Vári, L. and Kroon, D. (1986) Southern African Lepidoptera. A series of cross-referenced indices. Pretoria, South Africa: Transvaal Museum Bookshop. Available at: https://www.cabdirect.org/cabdirect/abstract/19891131574 (Accessed: 6 September 2022).

Wahlberg, N. et al. (2013) ‘Timing and patterns in the taxonomic diversification of Lepidoptera (butterflies and moths)’, PloS one, 8(11), p. e80875. Available at: https://doi.org/10.1371/journal.pone.0080875.

Zahiri, Reza et al. (2011) ‘A new molecular phylogeny offers hope for a stable family level classification of the Noctuoidea (Lepidoptera).’, Zoologica Scripta, 40(2), pp. 158–173.

Zahiri, R. et al. (2011) ‘A new molecular phylogeny offers hope for a stable family-level classification of the Noctuoidea (Lepidoptera)’, Zoologica Scripta, 40(2), pp. 158–173.

Zahiri, R. et al. (2012) ‘Molecular phylogenetics of Erebidae (Lepidoptera, Noctuoidea)’, Systematic Entomology, 37(1), pp. 102–124. Available at: https://doi.org/10.1111/j.1365-3113.2011.00607.x.

Zilli, A. and Grishin, N.V. (2019) ‘Unveiling one of the rarest “butterflies” ever (Lepidoptera: Hesperiidae, Noctuidae)’, Systematic Entomology, 44(2), pp. 384–395. Available at: https://doi.org/10.1111/syen.12330.

