## Supplementary figures and images for "The phylogenetic placement of an enigmatic moth *Egybolis vaillantina* based on museomics"

### Figure S1

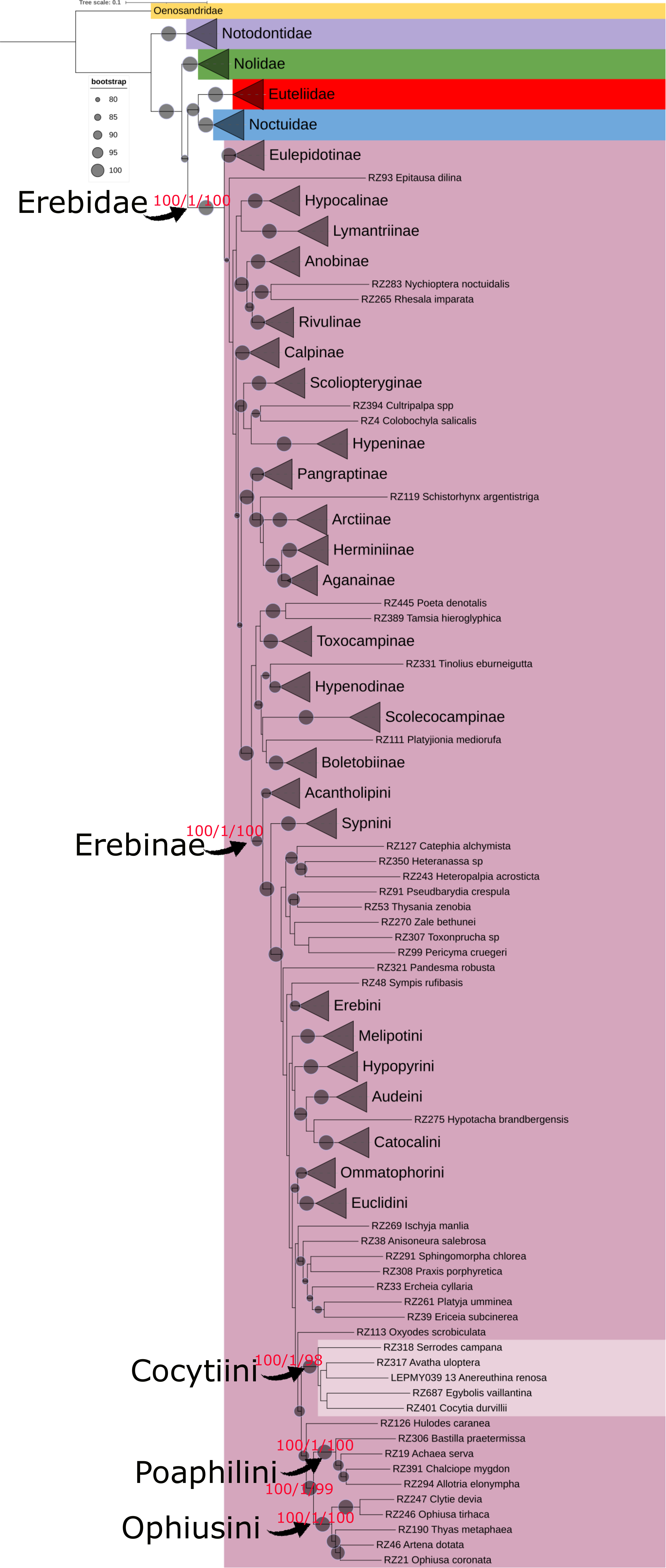
